# Differential contributions of an antimicrobial effector from *Verticillium dahliae* to virulence and tomato microbiota assembly across natural soils

**DOI:** 10.1101/2025.09.30.679524

**Authors:** Wilko Punt, Anton Kraege, Sabine Metzger, Natalie Schmitz, Jinyi Zhu, Stéphane Hacquard, Michael Bonkowski, Nick C. Snelders, Bart P.H.J. Thomma

## Abstract

Throughout their life cycle plants associate with diverse and complex microbial communities, known as their microbiota. These microbiota contribute to plant performance and health, in part by providing a microbial barrier against invading plant pathogens. To colonize plant hosts, pathogens not only have to overcome host immune responses, but also breach the microbial barrier, for which they secrete so-called effector proteins. Accordingly, the soil-borne fungal plant pathogen *Verticillium dahliae* secretes the antimicrobial effector Ave1 to suppress antagonistic microbes and facilitate host colonization. Notably, many pathogens, including *V. dahliae*, have life stages outside their host plants, for instance in soil, where they encounter diverse microbial communities. Yet, how antimicrobial effectors support establishment across these environments remains poorly understood. To address this, we established a collection of natural soil samples with diverse physicochemical properties and microbiota compositions. Using this collection, we show for three plant species, barley, tomato and cotton, that root-associated bacterial and fungal communities are primarily shaped by the type of soil, whereas the phyllosphere microbiota is mainly determined by plant species. On tomato, we furthermore show that Ave1 differentially contributes to virulence on diverse soils, as Ave1 altered the tomato microbiota on all soils tested, but the taxa affected by these shifts varied depending on the specific soils. Our findings suggest that while Ave1-mediated microbiota manipulation occurs across soils, its impact on fungal virulence is influenced by the specific composition of the soil-derived microbiota assembled by the host.

## INTRODUCTION

Plants host diverse microbial communities, known as the plant microbiota, which mainly include bacteria, fungi, and protists (Trivedi et al. 2020). These microorganisms colonize all plant parts, and together with the host plant, form a unified biological entity often referred to as the holobiont (Vandenkoornhuyse et al. 2015). Apart from seed-borne microbes inherited from the mother plant in the previous plant generation, the majority of microbes that make up the plant microbiota are recruited from environmental niches. While some microbes are transmitted through the air, the surrounding soil serves as the primary reservoir from which plants acquire most of their microbiota (Chialva et al. 2022). Soil properties such as pH, nutrient availability, organic carbon content, temperature and redox status shape the pool of microbes available for recruitment into the plant microbiota (Fierer, 2017). Consequently, the physicochemical properties of soil have a strong influence on plant microbiota assembly, as evidenced by the distinct microbial communities found in plants grown on different soils (Bulgarelli et al. 2012; Thiergart et al. 2020). At the same time, host genetics exert selective pressure on which taxa colonize and persist in the plant microbiota (Bulgarelli et al. 2012; Lundberg et al. 2012; Wagner et al. 2016). This is particularly evident in the formation of the core microbiota, which is a consistent subset of microbial taxa that reliably establish within the microbiota of a plant, even when plants are grown in diverse soils (Lundberg et al. 2012; Almario et al. 2022).

To date, numerous studies have separately demonstrated the importance of the bulk soil on the one hand, and of host genetics on the other hand, in structuring plant microbiota (Bulgarelli et al. 2012; Lundberg et al. 2012; Wagner et al. 2016; Fitzpatrick et al. 2018; Walters et al. 2018; Thiergart et al. 2020; Simonin et al. 2020; Tkacz et al. 2020). These studies have examined plant-associated microbes in diverse natural environments, where abiotic factors like local climate and weather can influence microbiota assembly, or have compared microbiota of different plant species grown in the same soil at a single location (Ofek-Lalzar et al. 2014; Wagner et al. 2016; Walters et al. 2018). However, studies that simultaneously evaluate the contribution of plant genetics and differential bulk soil microbiota on plant microbiota assembly, for instance by using various plant species in diverse natural soils while controlling for environmental influences, remain scarce (Tkacz et al. 2020, Dumack et al. 2022).

Microbes that establish in the plant microbiota interact with the host plant in various ways. Many microbes interact with plants as neutral commensals, while other microbes can be beneficial to the plant, or can be pathogenic and cause disease (Hassani et al. 2018). The community balance and composition of the microbiota plays an important role in plant health and performance, particularly by contributing to defense against pathogens (Du et al. 2025). Notably, plants have the ability to actively recruit beneficial microbes in response to pathogen attack. For instance, cucumber plants infected by the soil-borne pathogen *Fusarium oxysporum* f. sp. *cucumerinum* recruit *Bacillus amyloliquefaciens* to reduce disease severity (Liu et al. 2017). Over longer timescales, such plant-driven recruitment of beneficial microbes can result in the formation of disease-suppressive soils, where susceptible plants can grow in the presence of pathogens without experiencing severe disease symptoms (Du et al. 2025). A well-documented example, is the response of wheat plants to infection by *Gaeumannomyces graminis* var. *tritici*, the causal agent of “take-all” disease In this case, wheat recruits beneficial *Pseudomonas* species that antagonize the pathogen through the secretion of antimicrobial compounds, ultimately contributing to disease suppression over successive planting cycles in particular fields (Raaijmakers and Weller; Spooren et al. 2024). Importantly, protection via microbial recruitment is not limited to direct antagonism of pathogens. Some beneficial microbes enhance plant immunity through the induction of systemic defense responses (Pieterse et al. 2014). For example, *Arabidopsis thaliana* plants infected with the foliar pathogen *Hyaloperonospora arabidopsidis* (Hpa) selectively promote the growth of three bacterial species in the rhizosphere. This recruitment boosts systemic resistance to Hpa, improves overall plant growth, and can even benefit subsequent plant generations by fostering a protective microbiome (Berendsen et al. 2018). In this way, the plant microbiota has also often been considered as an additional layer of the immune system against pathogens by both inducing immune responses and directly antagonizing pathogens (Mendes *et al.* 2011; Carrión *et al.* 2019; Du et al. 2025; Durán et al. 2018).

While colonizing their hosts, plant pathogens secrete so-called effector molecules to promote host colonization by manipulating host physiology, including immunity (Jones and Dangl 2006; David E. Cook et al. 2015). Recently, several studies have demonstrated that pathogens exploit effector proteins that possess antimicrobial activity to manipulate the host microbiota, and thus facilitate colonization (Snelders et al. 2020; Chavarro-Carrero et al. 2024; Nick C. Snelders et al. 2021; Nick C. Snelders et al. 2023; Ökmen et al. 2023; Kettles et al. 2018; Chang et al. 2021; Gómez-Pérez et al. 2023; Mesny et al. 2024; Kraege et al. 2025). For example, the soil-borne fungal plant pathogen *Verticillium dahliae* exploits the antimicrobial effector protein Ave1 to suppress antagonistic Sphingomonadales bacteria during host colonization of tomato and cotton plants (Snelders et al. 2020). Interestingly, predictions from a machine learning tool suggest that 349 secreted *V. dahliae* effectors possess antimicrobial activity, indicating that *V. dahliae* may devote a substantial proportion of its secreted proteins to microbiota manipulation (Mesny and Thomma 2024).

Fungal pathogens such as *V. dahliae* occupy a range of ecologically distinct niches throughout their life cycle (Fradin and Thomma 2006; Guerreiro and Stukenbrock 2025). While they infect host plants during specific life stages, many also persist outside the host for extended periods, particularly in the soil (Fradin and Thomma 2006; Katan, 2017). Soil microbial communities are generally more diverse than those associated with plants and vary substantially depending on the physicochemical properties of the soil (Fierer 2017; Sokol et al. 2022). Accordingly, many pathogens are exposed to diverse microbial environments and must interact with a wide range of microbial taxa over time (Snelders et al. 2022). This is particularly relevant for broad host range pathogens like *V. dahliae*, which are adapted to numerous hosts and habitats and are thought to rely on antimicrobial effectors that facilitate interactions with different microbial communities (Trivedi et al. 2020; Snelders et al. 2022). Building on previous studies that explored antimicrobial effector functions using a single type of soil (Snelders et al. 2020), we hypothesize that the virulence contribution of antimicrobial effectors like Ave1, as well as their impact on microbial communities, may vary depending on the host-associated microbiota, which is largely determined by the bulk soil microbial community.

Here we report the establishment of a collection of natural soils that are diverse in both physicochemical characteristics and microbiota composition. We use this resource to simultaneously assess the contributions of the diverse types of soil and the plant genotype to plant microbiota assembly under controlled greenhouse conditions by analyzing fungal and bacterial communities associated with barley, cotton and tomato plants grown on each soil. Additionally, we utilize the soil collection to investigate the impact of the antimicrobial effector protein Ave1 on tomato microbiota composition and its role in *V. dahliae* virulence during infection of tomato plants harboring distinct microbiota.

## RESULTS

### Composing a collection of diverse natural soil samples

To study microbiota assemblies and the role of antimicrobial effector proteins of fungal plant pathogens in diverse soils we composed a collection of natural soil samples. We collected our soil samples in the Netherlands given the well-documented types of soil and the opportunity to sample a wide range of distinct types of soil on a relatively short geographical distance (Hartemink and Sonneveld 2013; Figure 1a). In total we collected samples from nine different natural soils which can be classified into the five major types of soil: river clay, sea clay, sand, peat and loam (Suppl. Table 1). Sampling sites were selected to avoid agricultural usage. In order to eliminate weeds and the majority of roots, the top 10 cm of soil was removed and the subsequent 30 cm of soil was collected. Besides the nine Dutch soils, we included the well-characterized and intensively studied Cologne agricultural soil (Bulgarelli et al. 2012).

**Figure 1.**
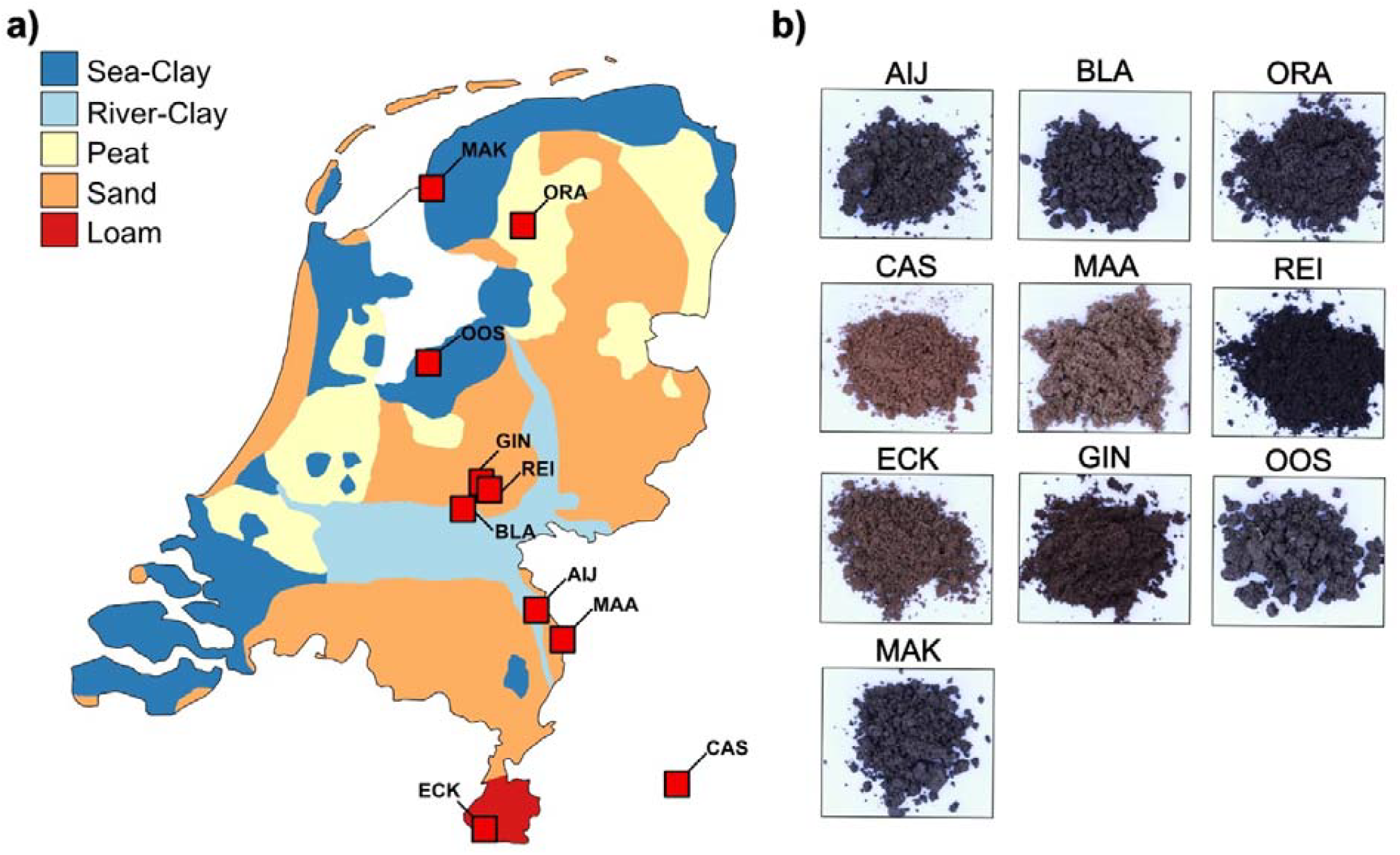
Establishment of a natural soil collection. **a)** Soil collection sites in the Netherlands. The map is colored according to major types of soil in the Netherlands. Sampling locations are indicated by red squares. **b)** Pictures of each soil from the soil sample collection.

The diversity of our soil sample collection is apparent from visible differences in soil texture and appearance (Figure 1b). To determine differences in physicochemical properties of our soil samples, we measured pH, the amount of total organic carbon and nitrogen, as well as element levels for all soil samples. The sandy soils (sand, peat, loam) displayed relatively low pH values, between 4.0 and 5.6, while the clay soils (river clay, sea clay) displayed higher pH values ranging from 6.2 to 7.7 (Figure 2a). With respect to carbon content, particularly the two river clay soils collected in Aijen (AIJ) and Blauwe Kamer (BLA) stood out with the highest carbon content of 4,83% and 7,01% respectively. The lowest carbon content was measured for the Cologne agricultural soil (CAS) with 0,26% and the sand soil collected in Maasduinen (MAA) with 0,21% (Figure 2b). A similar pattern was observed for the nitrogen content, as the highest value was measured for the river clay BLA with 0,35%, while lowest values were again determined for MAA at 0,006% and CAS at 0,02% (Figure 2C). Further, we also performed a total element analysis by conducting a HNO_3_-based element extraction followed by inductively coupled plasma mass spectrometry (ICP-MS) measurement. The elemental profiles of our soil samples were dominated by iron, calcium and aluminum (Figure 2d). Notably, when computing a principal component analysis (PCA) of the ICP-MS elemental raw data we observed separation according to types of soil, as the clay soils separated from the sandy soils and the CAS-soil (Figure 2e).

**Figure 2.**
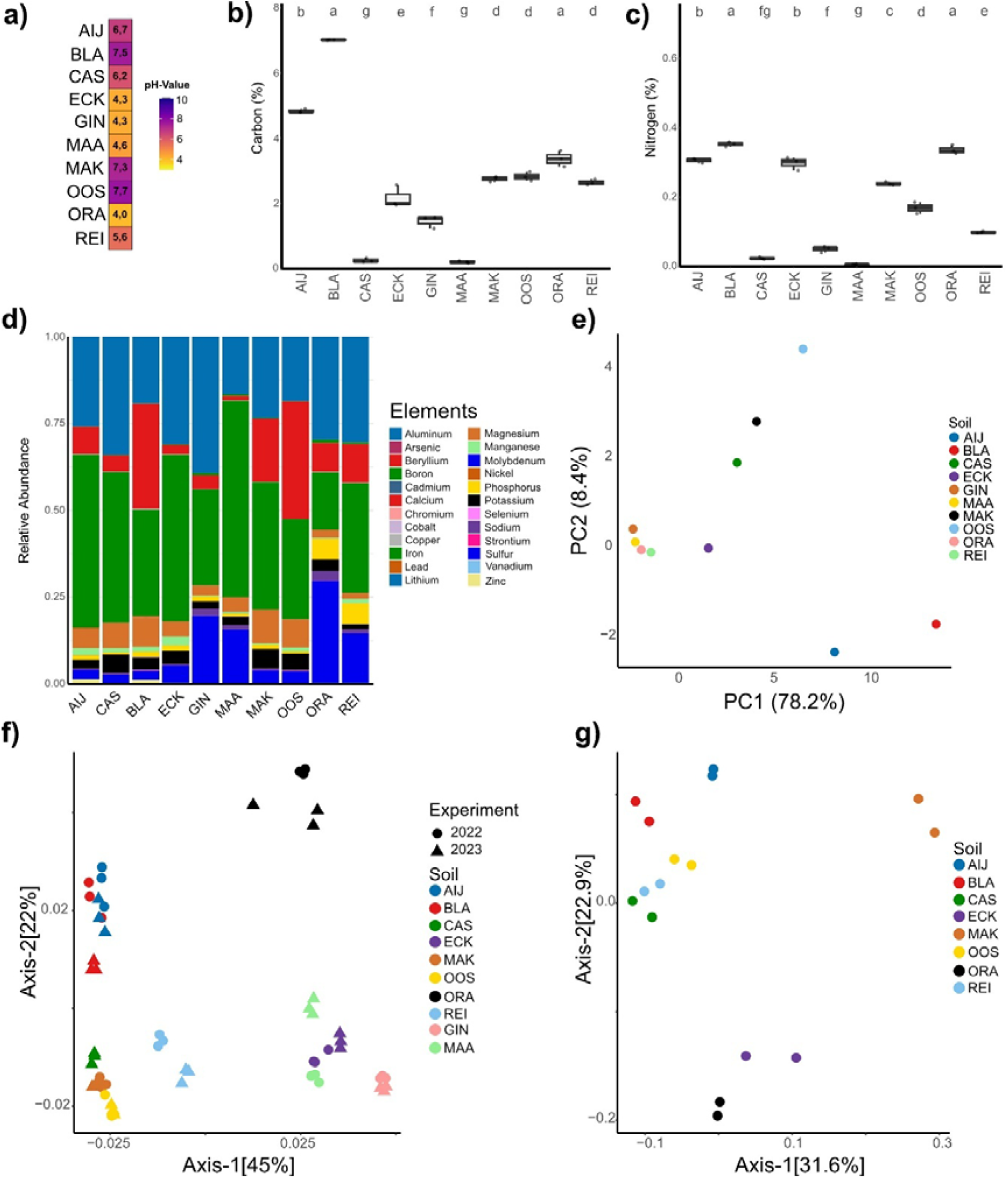
Physicochemical and microbiota analysis of the natural soil collection. **a)** Heatmap of pH-values. **b)** Boxplots displaying soil carbon contents. Different letters indicate statistical differences based on One-Way-Anova (Tukey HSD-Test pval < 0.05). **c)** Boxplots displaying soil nitrogen contents. Different letters indicate statistical differences based on One-Way-Anova (Tukey HSD-Test pval < 0.05). **d)** Relative abundance barplot for elements measured with ICP-MS. **e)** Principal component analysis (PCA) of the elemental profiles measured with ICP-MS. **f)** Principal coordinate analysis (PCoA) using weighted unifrac distances displaying bacterial bulk soil microbiota. Datapoints are shaped according to collection timepoint. **g)** PCoA using weighted unifrac distances displaying fungal bulk soil microbiota.

Many of the physicochemical properties are known to influence soil microbiota composition (Fierer 2017). To determine the bulk soil microbiota, we conducted 16S amplicon sequencing and analyzed the β-diversity by computing a principal coordinate analysis (PCoA) using the weighted Unifrac distance (Figure 2f). As expected, we observed separation of the microbiota according to the type of soil. Notably, we observed that apart from Reijerscamp (REI) the sandy soils collected from de Ginkelse Heide (GIN), Maasduinen (MAA), Oranjewoud (ORA) and ECK separate from the clay soil samples.

To investigate the consistency of the bulk soil microbiota, we compared the bulk soil microbiota of soil samples that were collected in two consecutive years; 2022 and 2023. In the PCoA, soils collected in the different years clustered, demonstrating a high degree of stability of these natural bulk soil microbiota (Figure 2f). Collectively, our data characterize the diversity of our natural soil sample collection with respect to physicochemical properties and bulk soil microbiota.

### Drivers of bacterial community assembly in roots and phyllosphere microbiota

Several studies have demonstrated that the soil as well as plant genetics influence plant microbiota assemblies (Bulgarelli et al. 2012; Lundberg et al. 2012; Wagner et al. 2016; Fitzpatrick et al. 2018; Walters et al. 2018; Thiergart et al. 2020; Simonin et al. 2020; Tkacz et al. 2020). These investigations typically involved plants collected from diverse natural environments, where microbiota assembly may additionally be affected by various abiotic factors, such as local climate and weather conditions, or they involve different plant species grown in the same soil at the same site (Ofek-Lalzar et al. 2014; Wagner et al. 2016; Walters et al. 2018). However, studies that simultaneously assess the contributions of different soils and of the plant genetics to microbiota assembly by examining diverse plant species grown in diverse natural soils while eliminating the impact of environmental factors remain scarce. Thus, we used our soil sample collection to investigate how plant-associated microbiota assemble across different plant species when grown under controlled conditions in a greenhouse. Specifically, we grew tomato (*Solanum lycopersicum*), cotton (*Gossypium hirsutum*), and barley (*Hordeum vulgare*) on the ten soils of our soil sample collection.

We first assessed how the diverse properties of the natural soils influence plant growth, by measuring plant canopy areas at three weeks after sowing. Cotton, tomato and barley plants grew on all soils except on the GIN and MAA soil samples, while tomato additionally failed to grow on ECK. Significant growth differences were observed across soils for each plant species (Suppl. Figure 2). Generally, the highest plant growth was observed on clay soil. For cotton the highest plant growth was determined on the MAK soil samples with an average canopy area of 39,76 cm^2^. Barley and tomato plants displayed highest plant growth on the BLA soil with barley plants reaching an average canopy area of 10,89 cm^2^ and tomato 22,23 cm^2^. Lowest plant growth for all three plants species was observed on the ORA soil, with average canopy areas of 23,44 cm^2^ for cotton, 1,8 cm^2^ for tomato plants and 1,42 cm^2^ for barley plants (Suppl. Figure 2). These results highlight the influence of the different soils on plant growth.

Next, we assessed the bacterial root and phyllosphere microbiota of the diverse plants grown on the soil collection by performing 16S rRNA sequencing. Bacterial communities in the root-associated microbiota were dominated by Proteobacteria, Actinobacteria, Acidobacteria, and Bacteroidetes across all soils and plant species. Notably, we observed considerable variation among individual plants of the same species grown in the same soil, despite prior homogenization. This may result from heterogeneity that persists in the natural soils samples even after mixing (Suppl. Figure 3). Nevertheless, as expected, we observed strong differences in bacterial community composition between plant species grown on the same soils. For instance, on the river clay soil AIJ, over 50% of the bacterial community in the barley root microbiota consisted of Proteobacteria, compared to only 25% of Proteobacteria in the tomato root microbiota. Rather, the tomato root microbiota on AIJ harbored higher proportions of Acidobacteria and Actinobacteria (Figure 3a). To assess the diversity of the root-associated microbial communities, we investigated microbial alpha diversities by calculating the Shannon index for each bacterial community sample. Notably, Shannon indices for the root microbiota varied across plant species and soils, with no plant species consistently exhibiting higher or lower diversity compared to the other species across the soils (Figure 3c). This was also supported by the calculated Hill numbers (Suppl. Figure 7). The lowest Shannon index was measured for cotton plants grown on ECK (2,49), whereas the most diverse communities were assembled by tomato plants grown on MAK (6,57).

**Figure 3.**
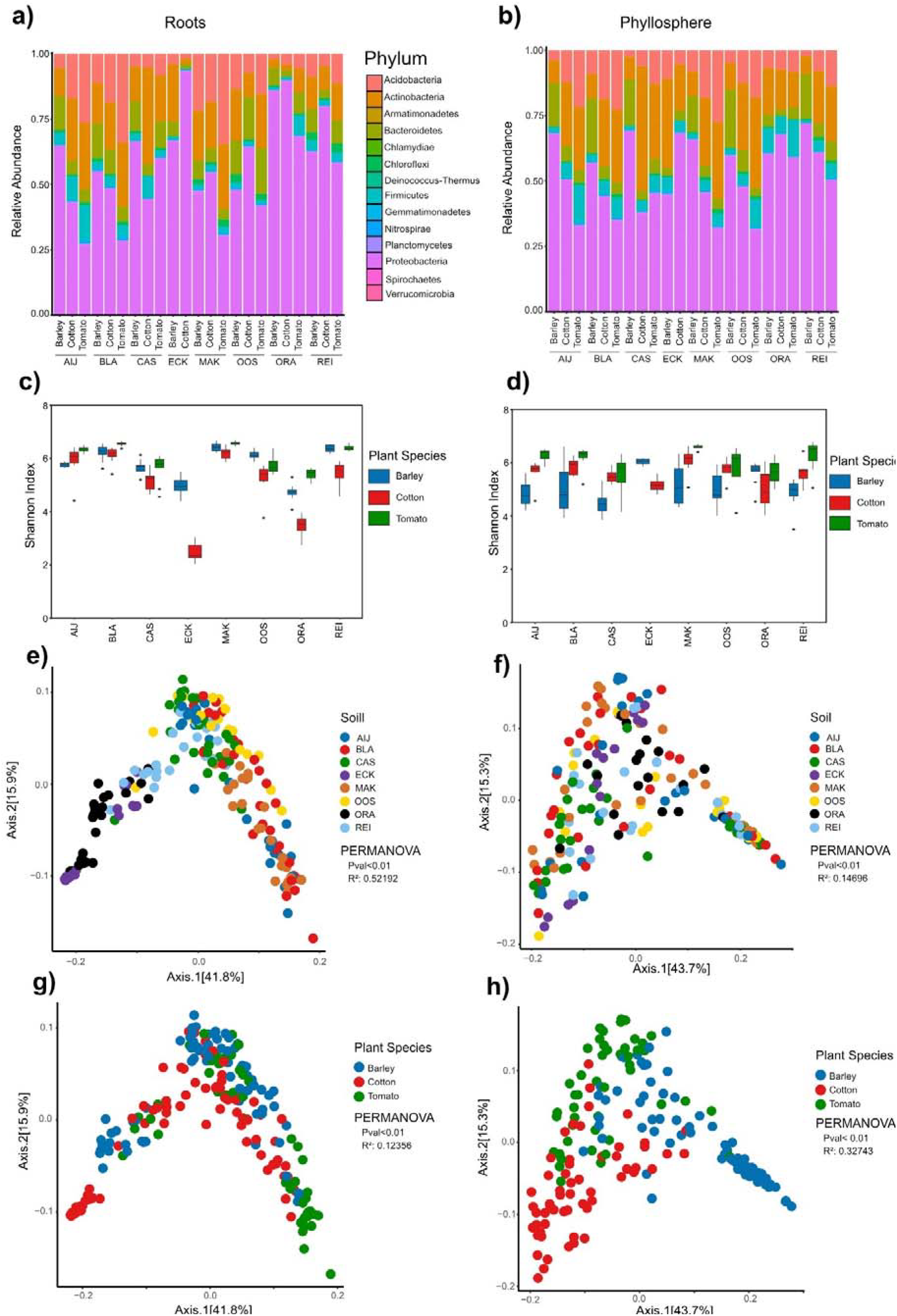
Bacterial composition of root and phyllosphere associated microbiota of barley, cotton and tomato plants grown on the different natural soils. **a)** Relative abundance in percentage on phylum level of the bacterial root microbiota. **b)** Relative abundance in percentage on phylum level of the bacterial phyllosphere microbiota. **c)** Shannon index of root microbiota. **d)** Shannon index of phyllosphere microbiota. **e)** Principal coordinate analysis (PCoA) based on weighted Unifrac distance of root microbiota. **f)** PCoA based on weighted Unifrac distance of phyllosphere. **g)** PCoA based on weighted Unifrac distance of root microbiota. **h)** PCoA based on weighted Unifrac distance of phyllosphere microbiota. All PERMANOVAs are performed with 9999 permutations.

To further disentangle the contributions of the type of soil and plant species to microbial diversity in the root microbiota, we analyzed β-diversities by conducting Principal Coordinate Analyses (PCoAs) based on weighted UniFrac distances. In the root-associated microbiota, bacterial communities grouped primarily according to thetype of soil, with sandy soils (ORA, ECK, REI) separating from clay soils (AIJ, BLA, OOS, MAK, CAS). The type of soil accounted for 52,2% of the observed variation within the microbiota, suggesting a dominant contribution to shaping root-associated bacterial communities (Figure 3e). Since we observed quite a “horseshoe effect” (Morton et al. 2017) we additionally calculated NMDS plots, which show a similar pattern (Suppl. Figure 8). Although also plant species significantly contributed to root-associated microbiota differentiation, it explained only 12,4% of the variation (Figure 3 g; Suppl. Figure 4b). Thus, root-associated microbiota are primarily structured according to the type of soil, and furthermore by plant species.

Next, we assessed if the patterns observed for root microbiota similarly hold true for phyllosphere microbiota. Similar to root microbiota, phyllosphere microbiota were dominated by Proteobacteria, Actinobacteria, Acidobacteria, and Bacteroidetes across all soils and plant species (Figure 3b). Also, for the phyllosphere microbiota we observed considerably variation between individual plants of the same species when grown in the same soil (Suppl. Figure 4). Notably, we observed strong differences between phyllosphere microbiota of different plant species grown in the same soil. Interestingly, these differences were similar across soils. For example, the tomato phyllosphere microbiota consistently exhibiting the lowest levels of Acidobacteria, followed by cotton and then barley in seven of the eight soils tested, with ECK as exception (Figure 3b). Next, we assessed community diversity in the phyllosphere microbiota by calculating Shannon indices. In the phyllosphere, the lowest Shannon indices were determined for barley plants grown on CAS (4,51) and AIJ (4,84), whereas highest values were again observed for tomato plants grown on MAK (6,59) and REI (6,27). Notably, the alpha diversity of bacterial phyllosphere microbiota displayed a more structured pattern when compared with the alpha diversity in the root microbiota, as barley consistently exhibited the lowest alpha diversity across six out of the eight soil samples, followed by cotton and then tomato (Figure 3d). This suggests that the plant species has a more pronounced influence on community diversity in the phyllosphere microbiota when compared with root-associated microbiota. We also analyzed β-diversities by conducting Principal Coordinate Analyses (PCoAs) based on the weighted UniFrac distances of the phyllosphere microbiota. Like the root-associated microbiota, the phyllosphere microbiota exhibited significant separation based on the type of soil, albeit that this explained substantially less variation (14,7%). Rather, plant species was the strongest determinant of the phyllosphere community composition, accounting for approximately 32,7% of the observed variation (Figure 3f, Suppl. Figure 3).

Collectively, our findings indicate that the soil is the strongest driver of bacterial microbiota diversity in the root microbiota, while plant species plays a more significant role in shaping bacterial phyllosphere communities.

### Drivers of fungal community assembly in root-associated and phyllosphere microbiota

To assess whether patterns observed for bacterial microbiota across plant species grown on our soil collection also apply to the fungal component of the microbiota, we conducted ITS sequencing. First, we examined the fungal communities in the bulk soil microbiota of the eight soil samples used for the plant microbiota assembly study. This analysis revealed that the sand-like soils ECK and ORA separate from the clay soil. The REI soil, although also a sandy-soil, grouped with the clays. This indicates that the soil sample collection harbors distinct fungal communities (Figure 2g).

Analysis of the fungal communities in the root-associated microbiota revealed that fungal communities across plant species and type of soil were dominated by fungal species from the phyla Ascomycota, with Basidiomycota and Mortierellomycota (Figure 4a). Notably, the fungal composition of the root microbiota is also influenced by plant species across soil samples. For instance, on ECK, the fungal communities in the barley root microbiota contained more than 80% Ascomycota, while the fungal root microbiota of cotton plants contained only 50% Ascomycota, with a substantially higher abundance of Basidiomycota (Figure 4a; Suppl. Figure 6). Shannon index calculations revealed lower alpha diversities of the root-associated fungal communities when compared with bacterial communities, with no consistent patterns of alpha diversity based on the plant species emerging across soil samples. Analysis of the β-diversity by performing a PCoA using the weighted Unifrac distance matrix revealed that root-associated fungal communities separate based on the soil sample in which the plants were grown, explaining 31% of the variation observed in the fungal microbiota (Figure 4e). Root-associated fungal communities also displayed weak separation according to plant species, which explained 9% of the variation (Figure 4g; Suppl. Figure 6b). Overall, these findings suggest that fungal communities in the root-associated microbiota are primarily shaped by the type of soil. As expected, also in the phyllosphere microbiota the fungal communities were dominated by Ascomycetes, followed by Basidiomycetes and Mortierellomycetes (Figure 4b; Suppl. Figure 5). Similar as for the alpha diversity in the root-associated fungal microbiome we did not observe any alpha diversity patterns based on plant species or the type of soil in the fungal phyllosphere microbiota (Figure 4d). The β-diversity analysis of the fungal community in the phyllosphere microbiota revealed weak separation based on the type of soil, which explained 13% of the variation (Figure 4f). Notably, similar as for the bacterial phyllosphere microbiota, we observed strong separation of the fungal phyllosphere community based on plant species, which explained 26% of the variation (Figure 4h; Suppl. Figure 5b). Collectively, our dataset reveals that fungal communities in the root-associated microbiota are more strongly influenced by types of soilthan by plant species, while the plant species acts as the primary driving factor for fungal communities in the phyllosphere microbiota.

**Figure 4.**
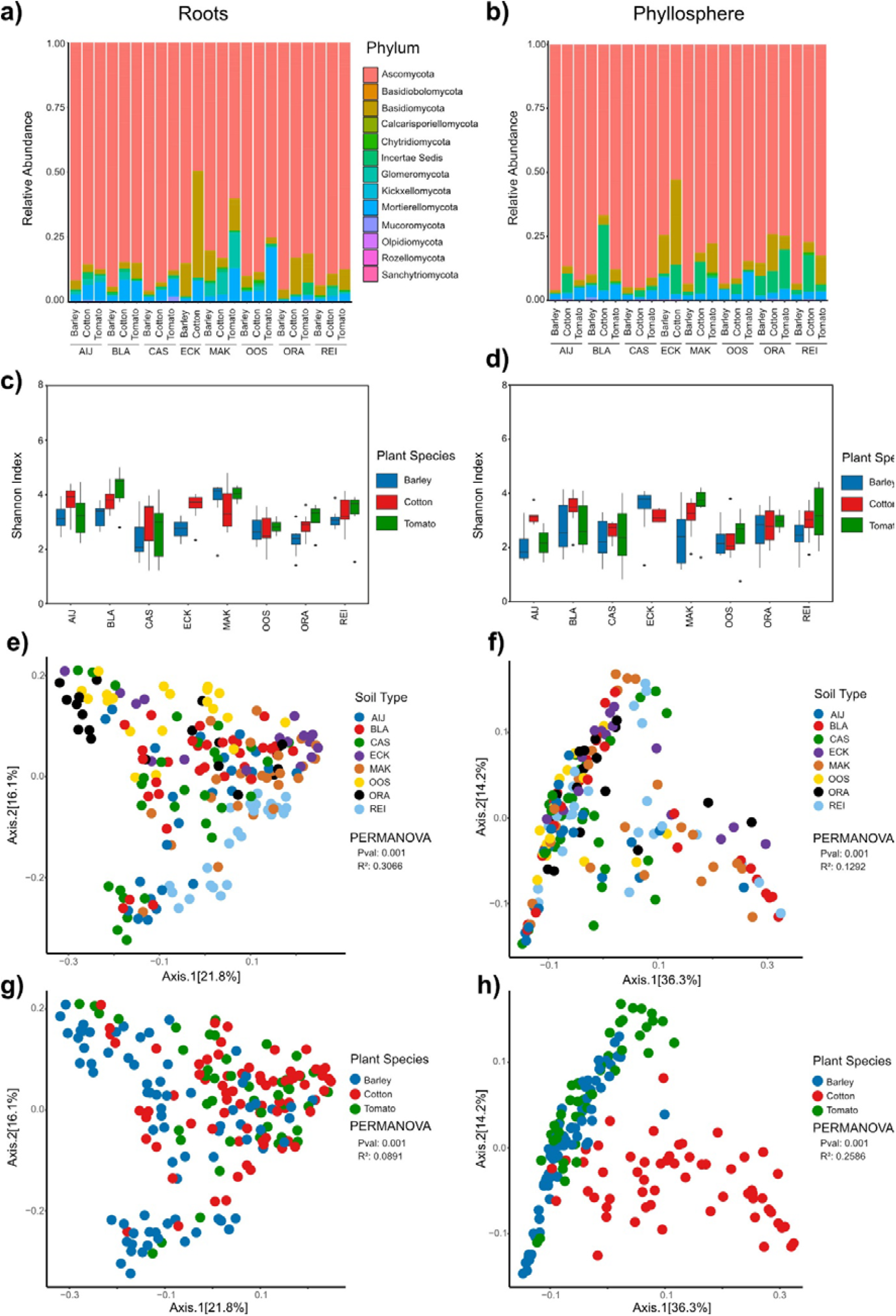
Composition of the fungal root and phyllosphere associated microbiota of barley, cotton and tomato plants grown on the different natural soils. **a)** Relative abundance in percentage on phylum level of the fungal root microbiota **b)** Relative abundance in percentage on phylum level of the fungal phyllosphere microbiota **c)** Shannon index of root microbiota. **d)** Shannon index of phyllosphere microbiota. **e)** Principal coordinates analysis based on weighted Unifrac distance of root microbiota. **f)** PCoA based on weighted Unifrac distance of phyllosphere **g)** PCoA based on weighted Unifrac distance of root microbiota **h)** PCoA based on weighted Unifrac distance of phyllosphere microbiota All PERMANOVAs are performed with 9,999 permutations.

### Differential contribution of antimicrobial effectors to fungal virulence across types of soil

The plant microbiota plays an important role in plant health, fitness and defense against plant pathogens (Trivedi et al. 2020). To colonize their hosts, plant pathogens have evolved antimicrobial effector proteins to manipulate host-associated microbiota (Mesny et al. 2024). For instance, *V. dahliae* uses the antimicrobial effector Ave1 to suppress antagonistic microbes during host colonization. Ave1 was demonstrated to facilitate host colonization of cotton and tomato plants grown in potting soil through targeting, amongst others, antagonistic Sphingomonadales bacteria (Snelders et al. 2020). As a globally distributed soil-borne pathogen with a broad host range, *Verticillium dahliae* successfully colonizes host plants across diverse types of soil, which likely harbor distinct microbial communities (Klimes et al. 2015; Singh et al. 2025). We hypothesized that the outcome of effector-mediated microbiota manipulation may vary depending on the host-associated microbiota, which is largely assembled from the surrounding bulk soil microbiota. To address this, we assessed the virulence contribution of the antimicrobial effector Ave1 by growing tomato plants on our soil collection and inoculating them with either wild-type *V. dahliae* or an *Ave1* deletion mutant (Jonge et al. 2012; Snelders et al. 2020). We observed a significant reduction in biomass of tomato plants inoculated with the wild type strain when compared with plants inoculated with the *Ave1 deletion st*rain on AIJ, BLA, ORA and MAK, indicating that Ave1 contributes to fungal virulence on these soils. In contrast, no such difference was observed for plants grown in OOS and REI, suggesting that Ave1 differentially contributes to fungal virulence across soils (Figure 5a). To rule out the possibility that the observed differences between soil types were due to coinfections with naturally occurring *Verticillium* strains, we quantified the relative abundance of *Verticillium* in bulk soil and found only low abundances with no significant differences (Suppl. Figure 9). Previous work demonstrated that Ave1 also negatively impacts the abundance of other taxa, including Verrucomicrobiales, Chitinophagaceae, Flavobacteriales and Burkholderiales during infections of cotton and tomato plants grown on potting soil (Snelders et al. 2020). We then asked whether variation in the abundance of these bacteria in the root-associated microbiota of tomato plants could explain the differences in virulence contribution of Ave1 across soils. To test this, we measured their relative abundance in tomato plants grown in the different natural soil samples. Of the tested taxa, Sphingomonadales, Flavobacteriales, and Burkholderiales showed no significant differences in relative abundances across soils. While significant variation in relative abundance was observed for the Verrucomicrobiales and Chitinophagaceae on several soils, these differences did not correlate with the observed Ave1-related virulence phenotype (Suppl. Figure 10a).

**Figure 5.**
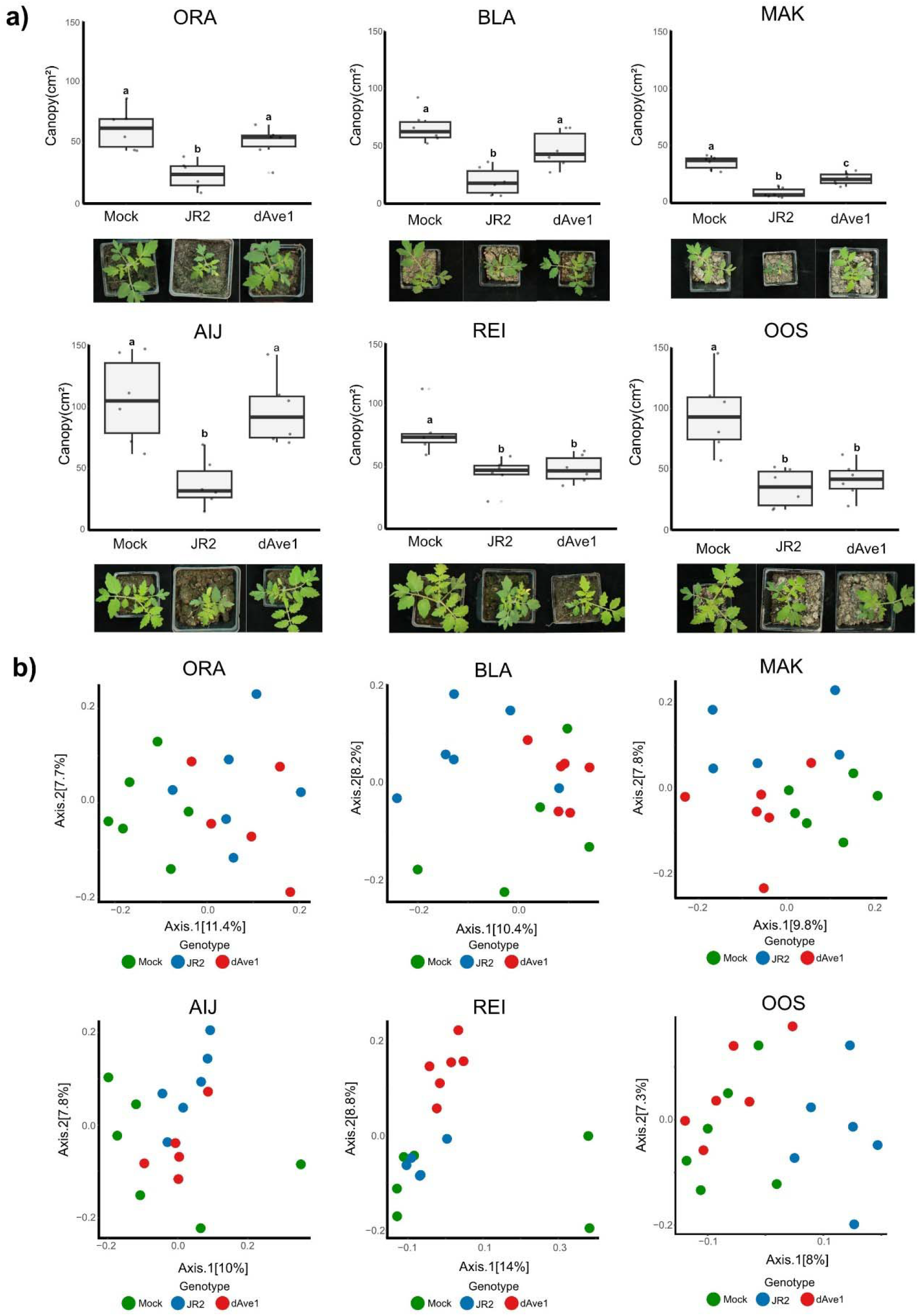
Antimicrobial effector Ave1 differentially contributes to virulence of *Verticillium dahliae* depending on the soil. **a)** Canopy area in cm^2^ of tomato plants grown on the different natural soils at 14 dpi with wild-type *V. dahliae* (JR2) or an Ave1 deletion mutant (dAve1). Different letters indicate statistical differences based on One-Way-Anova (Tukey HSD-Test pval < 0.05). Pictures display a representative plant per treatment. b) Principal coordinate analysis (PCoA) based on Unifrac distances of the root microbiota of tomato plants grown on different soils at 14 dpi with wild-type *V. dahliae* (JR2) or an *Ave1* deletion mutant (dAve1).

To assess the impact of Ave1 on the tomato root-associated microbiota we investigated the microbiota composition of tomato plants that were mock-inoculated, or inoculated with *V. dahliae* strain JR2 or the *Ave1* deletion mutant (Figure 5b). By computing a PCoA based on UniFrac distances we observed that the tomato microbiota from plants inoculated with the wild type and the deletion mutant consistently separated across all soils, except for the ORA soil (Suppl. Figure 11). Notably, we also observe such separation in the microbiota of plants grown on REI and OOS, even though we did not detect a virulence contribution of Ave1.

Further, to investigate the bacterial taxa affected by Ave1 on the natural soils we conducted differential abundance analysis between the microbiota of plants inoculated with *V. dahliae* strain JR2 or the *Ave1*-deletion mutant. This analysis revealed significant shifts in microbiota composition at the genus level across all soil samples, including OOS and REI, even though no virulence contribution of Ave1 was observed on these soils. Notably, on each of the soils the effector causes distinct shifts in the microbiota (Suppl. Figure 10b). Collectively, our data indicates that the outcome of effector-mediated microbiota manipulation by *V. dahliae* is determined by the composition of the host-associated microbiota which, in turn, is influenced by the surrounding soil.

## DISCUSSION

Plant microbiota contribute substantially to plant productivity, in part by serving as an additional barrier against invading pathogens (Mesny et al. 2024; Du et al. 2025). Over recent years, it has become evident that plant pathogens manipulate host microbiota through the secretion of antimicrobial effector proteins in turn, thus facilitating niche establishment and host colonization (Snelders et al. 2020; Chavarro-Carrero et al. 2024; Snelders et al. 2021; Snelders et al. 2023; Ökmen et al. 2023; Kettles et al. 2018; Chang et al. 2021; Gómez-Pérez et al. 2023; Mesny et al. 2024; Kraege et al. 2025). Notably, many pathogens spend parts of their life cycles outside their hosts, where they encounter diverse microbial communities. However, how antimicrobial effectors aid fungal establishment across these diverse environments is still poorly understood. Here, we present a collection of natural soils that we thoroughly characterized in terms of their physicochemical properties as well as their microbiota compositions. Using this soil collection, we reveal that the antimicrobial effector protein Ave1 from soil-borne fungal plant pathogen *Verticillium dahliae*, which was previously demonstrated to facilitate host colonization through the suppression of antagonistic Sphingomonadales bacteria (Snelders et al. 2020), *contributes* to fungal virulence on tomato plants only in a subset of these soils. Our finding suggests that the virulence contribution of this effector is determined by the soil on which the host plant grows. Interestingly, differential virulence contributions have similarly been reported for another antimicrobial effector from *V. dahliae*, called Av2. While initially no contribution to fungal virulence was recorded (Chavarro-Carrero et al. 2021), a subsequent study using a different growth substrate, likely with a distinct microbiota, revealed that Av2 interfered with the host plant’s ‘cry for help’ recruitment of beneficial *Pseudomonas* bacteria, leading to a clear virulence contribution of the effector (Kraege et al. 2025). These differences in virulence contributions of antimicrobial effectors are likely due to variation in soil microbiota, which impacts the composition of plant-associated microbial communities encountered by the pathogen during infection in turn. Interestingly, our microbiota analyses revealed that the Ave1 effector significantly altered the tomato microbiota on all tested soils. This implies that microbiota manipulation by the effector does not necessarily translate into measurable contributions to fungal virulence and thus, that this effector does not solely target antagonists of *V. dahliae* growth. We therefore infer that the presence or absence of antagonistic microbes that can be impacted by an antimicrobial effector will determine whether that effector contributes to fungal virulence during host infection. This hypothesis is supported by observations made for the *V. dahliae* antimicrobial effector protein Ave1L2. A previous study investigating Ave1L2 demonstrated that in communities artificially depleted of antagonistic Actinobacteria, described as a crucial target of the effector, the protein still impacted community composition albeit without a measurable virulence contribution (Snelders et al. 2023).

Notably, the observed impact that Ave1 caused on the plant microbiota substantially differed across soils. Many antimicrobial effector proteins do not specifically act on a single antagonistic microbe, but rather act on multiple plant microbiota members, thus exerting broader, system-level impacts on microbial communities (Snelders et al. 2020; Snelders et al. 2021; Chavarro-Carrero et al. 2024; Kraege et al. 2025). Since plant microbiota function as networks of interdependent species (van der Heijden and Hartmann 2016), changes affecting one member can cascade through the community. Thus, removal or suppression of particular microbes by fungal effectors may trigger cascading shifts in community structure and function due to these intermicrobial interactions. This interconnectedness implies that the effects of antimicrobial effector activity on the microbiota can vary substantially between microbial communities, driven by the unique web of intermicrobial interactions in each environment.

Our study additionally provides a controlled comparison of how both types of soil and plant genotype influence microbiota assembly across different plant compartments. While previous studies have independently demonstrated that rhizosphere communities are primarily shaped by soil and phyllosphere communities by host genotype (Bulgarelli et al. 2012; Lundberg et al. 2012; Wagner et al. 2016; Fitzpatrick et al. 2018; Walters et al. 2018; Thiergart et al. 2020; Simonin et al. 2020; Tkacz et al. 2020) these insights were often derived from field studies conducted in divergent natural environments, where additional abiotic factors such as climate and weather may influence microbiota composition, or from experiments that varied either soil or plant species, but rarely both. Tkacz et al. (2020) assessed microbiota assembly across four plant species grown in two distinct soils and demonstrated that soil has a stronger influence than plant species on shaping rhizosphere microbiota. In our study, we extend these findings by using a different set of plant species and a broader collection of ten diverse, well-characterized soils, including the Cologne agricultural soil (Bulgarelli et al. 2012; Yang Bai et al. 2015) and Reijerscamp soil (Berendsen et al. 2018; Poppeliers et al. 2024) under highly controlled greenhouse conditions. We not only confirm that the type of soil plays a dominant role in rhizosphere microbiota assembly, but also show simultaneously that, in contrast, phyllosphere communities are primarily shaped by plant species rather than the type of soil. Notably, this work also demonstrates a rhizosphere effect on fungi, an important aspect of microbiome assembly that is underexplored when compared with the effect on bacteria.

Taken together, our findings support the view that antimicrobial effector proteins are context-dependent components of fungal secretomes, rather than universally acting virulence factors with consistent effects across environments. Notably, a recent machine-learning analysis predicted that, for several fungi, at least one-third of effector proteins possess antimicrobial activity (Mesny and Thomma 2024), suggesting that fungi may deploy large repertoires of such antimicrobial effectors to establish themselves in diverse environments. Deeper insight into their functions and the mechanisms underlying this environmental variability will not only advance our understanding of fungal niche adaptation but may also inform the development of more robust, microbiota-based disease control strategies for agriculture.

## MATERIALS AND METHODS

### Soil collection and storage

Natural soils were collected Three soil collections were performed, in January 2022, February 2023 and February 2024. at nine sites in the Netherlands: Makkum (53°05’09.8”N 5°26’20.3”E), Oranjewoud (52°57’11.7”N 5°57’45.6”E), Ginkelse Heide (52°02’10.7”N 5°43’38.9”E), Eckelrade (50°47’57.7”N 5°44’42.5”E) Maasduinen (51°28’34.3”N 6°11’34.9”E), Oostvaardersplassen (52°27’50.0”N 5°25’10.8”E), Reijerscamp (52°00’37.7”N 5°46’25.0”E), Blauwe Kamer (51°56’34.4”N 5°37’12.9”), Aijen (51°34’55.0”N 6°02’27.3”E)in three consecutive years in February from 2022-2024. For collection, the top 10 cm of soil was removed and the subsequent 30 cm of soil was collected. After collection, soil samples were homogenized and rocks and pieces of plant material were removed before the soil was stored in sealed buckets at 8℃ until further use. Further, Cologne agricultural soil (50°57’27.8”N 6°51’22.4”E; Bai *et al*., 2015) was included.

### Physicochemical soil analysis

For physicochemical analysis, 50 g of each soil was freeze dried and ground to fine powder using a mortar and pestle. To measure soil pH, ground soil powder was suspended with 150 ml of distilled water and incubated for 1 hour. Subsequently the pH was measured using a pH-electrode (Meddler Toledo, Giessen, Germany). Carbon and nitrogen levels were measured using the FLASH2000 CHNS/O analyzer (Thermo Fisher Scientific, Waltham, USA). To measure elemental contents, 100 mg of soil powder was weighed into metal-free centrifugation tubes (VWR, Radnor, USA). Samples were the soaked in 500 µl of 30% nitric acid for 2 hours. Subsequently, the volume was adjusted to 1 ml with 30% nitric acid and the sample was incubated for 14 hours at 65℃. Next, the suspension was incubated at 95℃ for 90 minutes. Samples were cooled to room temperature and 200 µl of hydrogen peroxide were added. Subsequently, the samples were incubated at 95℃ for 30 minutes. Next, the samples were diluted to 10 ml using MQ-water and centrifuged at 13,000 rpm for 1 hour at 4℃. The supernatant was transferred to a clean metal-free 50 ml centrifugation tube and incubated at 4℃ overnight, followed by centrifugation at 13,000 rpm for 1 hour at 4℃. Finally, 600 µl of supernatant were mixed with 2,4 ml of 2% nitric acid. ICP-MS measurements were carried out on an Agilent 7700 ICP-MS (Agilent Technologies, Waldbronn, Germany) in the Biocenter MS-Platform of the University of Cologne. All measurements were performed in technical triplicates and strictly followed the manufacturer`s instructions using He in the collision cell mode to minimize spectral interference.

### Plant growth assays

Tomato (*Solanum lycopersicum L.*) cultivar MoneyMaker, barley (*Hordeum vulgare*) cultivar GoldenPromise, Cotton (*Gossypium hirsutum*) cultivar DDHY642201-AC and Arabidopsis (*Arabidopsis thaliana*) ecotype Col-0 were used for all assays. Before sowing, seeds were surface-sterilized using chlorine gas generated by adding 3 mL of hydrochloric acid (HCl) to 100 mL of bleach (sodium hypochlorite) in a 250 mL beaker placed inside a glass container sealed with a lid and parafilm and incubated for 5 hours. After sterilization, the container was vented in a fume hood overnight. Subsequently, surface sterilized seeds were sown on soil and grown for three weeks in a greenhouse chamber with 16 hours of light at 23°C, followed by 8 hours in darkness at 22°C. Plant growth was assessed by calculating canopy areas, for tomato and cotton based on overhead pictures and for barley plants based on side pictures using ImageJ (Schneider *et al*., 2012). Subsequently, plants were harvested for microbiota analysis. Tomato and cotton phyllosphere samples were collected by harvesting the stem from the soil-line to the cotyledons, while barley phyllosphere samples were collected by harvesting the first 5 cm of plant tissue above the soil line. To collect root microbiota samples, plants were uprooted and loose soil was removed from the root system through gentle shaking.

### Microbiota sequencing

Samples were manually ground to fine powder using a mortar and pestle. Subsequently, 400 mg of tissue or soil were used for DNA extraction using the DNeasy PowerSoil Pro Kit (Qiagen, Venlo, The Netherlands). Next, DNA was further purified using the Monarch PCR&DNA Clean Up kit (New England Biolabs, Ipswich, USA). DNA purity and concentration were assessed using the Qubit 4 fluorometer (Thermo Fisher Scientific, Waltham, USA) and the Nanodrop 2000 spectrophotometer (Thermo Fisher Scientific, Waltham, USA). DNA was used for the amplification of the variable regions 3-4 of the 16S region using primers 341f (ACTCCTACGGGAGGCAGCAG) and 806r (GGACTACHVGGGTWTCTAAT) in the presence of the mPNA (GGCAAGTGTTCTTCGGA) and pPNA (GGCTCAACCCTGGACAG) blocking clamps (PNABio, Newbury Park, USA). Additionally, amplification of the ITS2 region was conducted using the primers ITS3 (GCATCGATGAAGAACGCAGC) and ITS4 (TCCTCCGCTTATTGATATGC) in the presence of the ITS2 PNA (CGAGGGCACGTCTGCCTGG) blocking clamp (PNABio, Newbury Park, USA). All amplicons were sequenced on an Illumina MiSeq Platform (BGI-Genomics, Shenzhen, China). For the bulk soil microbiota from the 2022 and 2023 collections, the V5-V7 regions were amplified with primers 799F (AACMGGATTAGATACCCKG) and 1139R (ACGTCATCCCCACCTTCC) and amplicons were similarly sequenced on an Illumina Miseq Platform (Cologne Center for Genomics, Cologne, Germany). Only samples with at least 10.000 reads were considered for the analysis. Data analysis was conducted as described previously (Callahan *et al*., 2016; Snelders *et al*., 2020).

### Microbiota analysis

Sequencing data were processed using R v.4.2.0. as described previously (Callahan et al., 2016; Snelders et al., 2020). In brief, reads were demultiplexed with cutadapt (v4.1; Martin, 2011), then trimmed and filtered to an average paired read length of 412 bp with a Phred score of 30. OTUs were inferred from the trimmed reads using the DADA2 method (v 1.24; Callahan et al., 2016). Taxonomy was assigned using the Ribosomal Database Project (RDP,v 18; Cole et al., 2014). The pyloseq package (v1.40.0; McMurdie & Holmes, 2013) was used to calculate α- and β-diversity, while PERMANOVA was conducted with the vegan package (v2.6-4; Oksanen et al., 2004) package. Differential abundance analysis was done using the DESeq2 package (v1.36.0; Love et al., 2014) using a negative binomial Wald test and a significance P adjusted threshold < 0.05.

### *Verticillium* inoculation assays

*Verticillium dahliae* inoculations were conducted on 10-day-old tomato plants. Inoculum was prepared by harvesting conidiospores of 10-day-old cultures of *V. dahliae* strain JR2 and an *Ave1* deletion mutant (de Jonge et al., 2012; Snelders *et al*., 2020) on potato dextrose agar (PDA; Carl Roth, Karlsruhe, Germany). The collected conidiospores were washed three times in MQ water, each time followed by centrifugation at 10.000 rpm for 10 minutes. Subsequently, the conidiospores were counted using a Neubauer chamber and the inoculum concentration was adjusted to 10^6^ conidiospores/ml. For the inoculations, plants were uprooted and the roots were rinsed with MQ-water before being placed into the conidiospore suspension for 8 minutes. Subsequently, plants were planted back into the soil. Disease symptoms were monitored at 14 dpi by measuring the tomato canopy area based on overhead pictures using ImageJ (Schneider *et al*., 2012).

## AUTHOR CONTRIBUTIONS

W.P., A.K., N.C.S and B.P.H.J.T. conceived the project. W.P., A.K., N.C.S and B.P.H.J.T. designed the experiments. S.H. provided biological materials. W.P., A.K., J.Z., S.M., M.B., N.S. and N.C.S. performed the experiments. W.P., A.K., N.C.S and B.P.H.J.T. analyzed the data. W.P., A.K. and B.P.H.J.T. wrote the manuscript. All authors read and approved the final manuscript.

## ACKNOWLEDGEMENTS

B.P.H.J.T. acknowledges funding by the Alexander von Humboldt Foundation in the framework of an Alexander von Humboldt Professorship endowed by the German Federal Ministry of Education and is furthermore supported by the Deutsche Forschungsgemeinschaft (DFG, German Research Foundation) under Germany’s Excellence Strategy – EXC 2048/1 – Project ID: 390686111.

## DATA AVAILABILITY

16S profiling data are available in the NCBI Genbank database under BioProject PRJEB95937.

