## Supplemental Information for "Differential contributions of an antimicrobial effector from *Verticillium dahliae* to virulence and tomato microbiota assembly across natural soils"

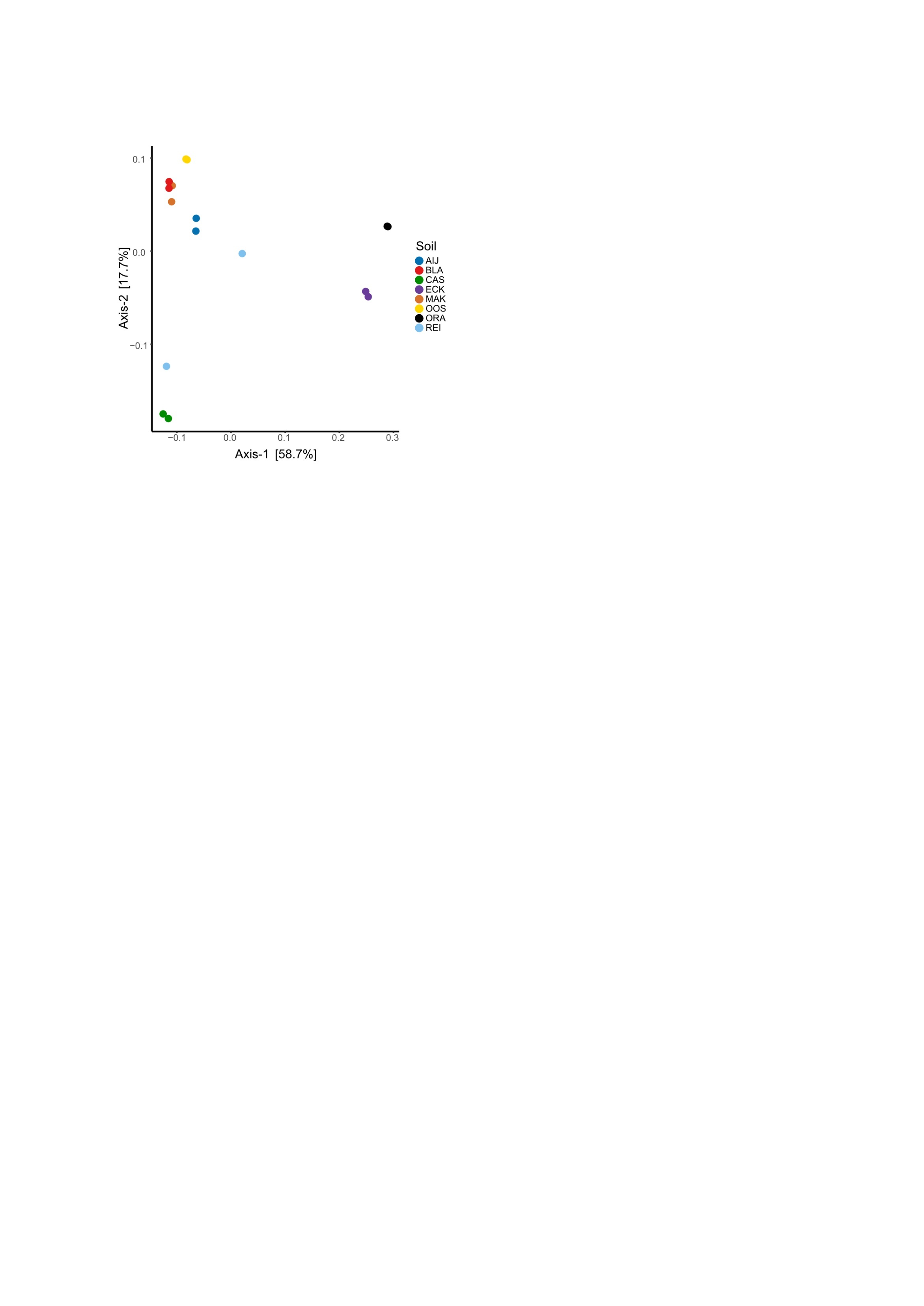
**Supplementary Figure 1. Principal coordinate analysis (PCoA) based on weighted unifrac distances of the bacterial bulk soil microbiota from soil samples collected in 2024.**

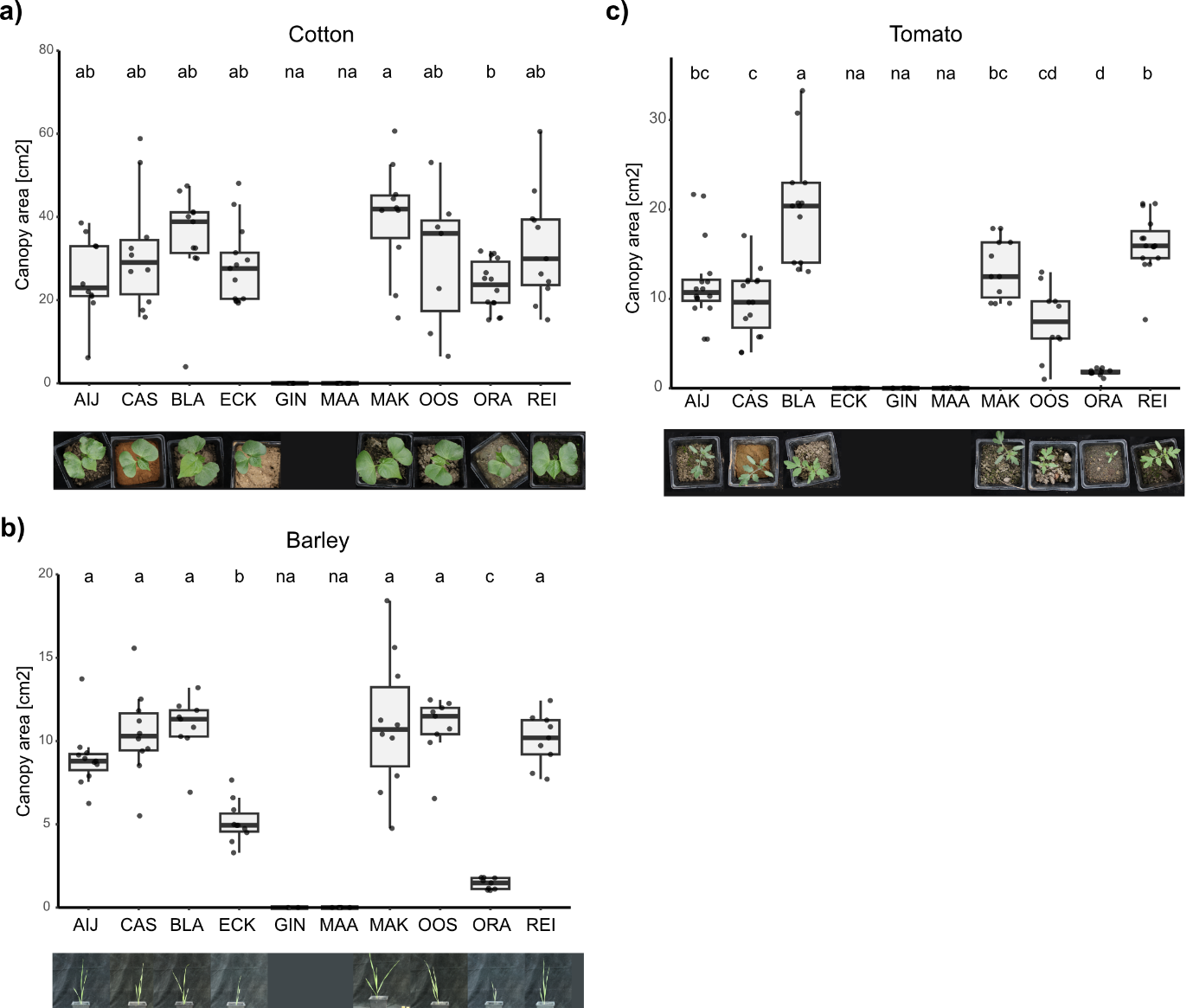
**Supplementary Figure 2. Plant growth on natural soils. a)** Canopy area of cotton plants grown on natural soils for 21 says. **b)** Canopy area of tomato plants grown on natural soils for 21 says. **c)** Canopy area of barley plants grown on natural soils for 21 says. Different letters indicate statistical differences based on One-Way-Anova (Tukey HSD-Test pval < 0.05) for each panel. Soils that did not support plant growth were excluded from the statistics and labeled na.

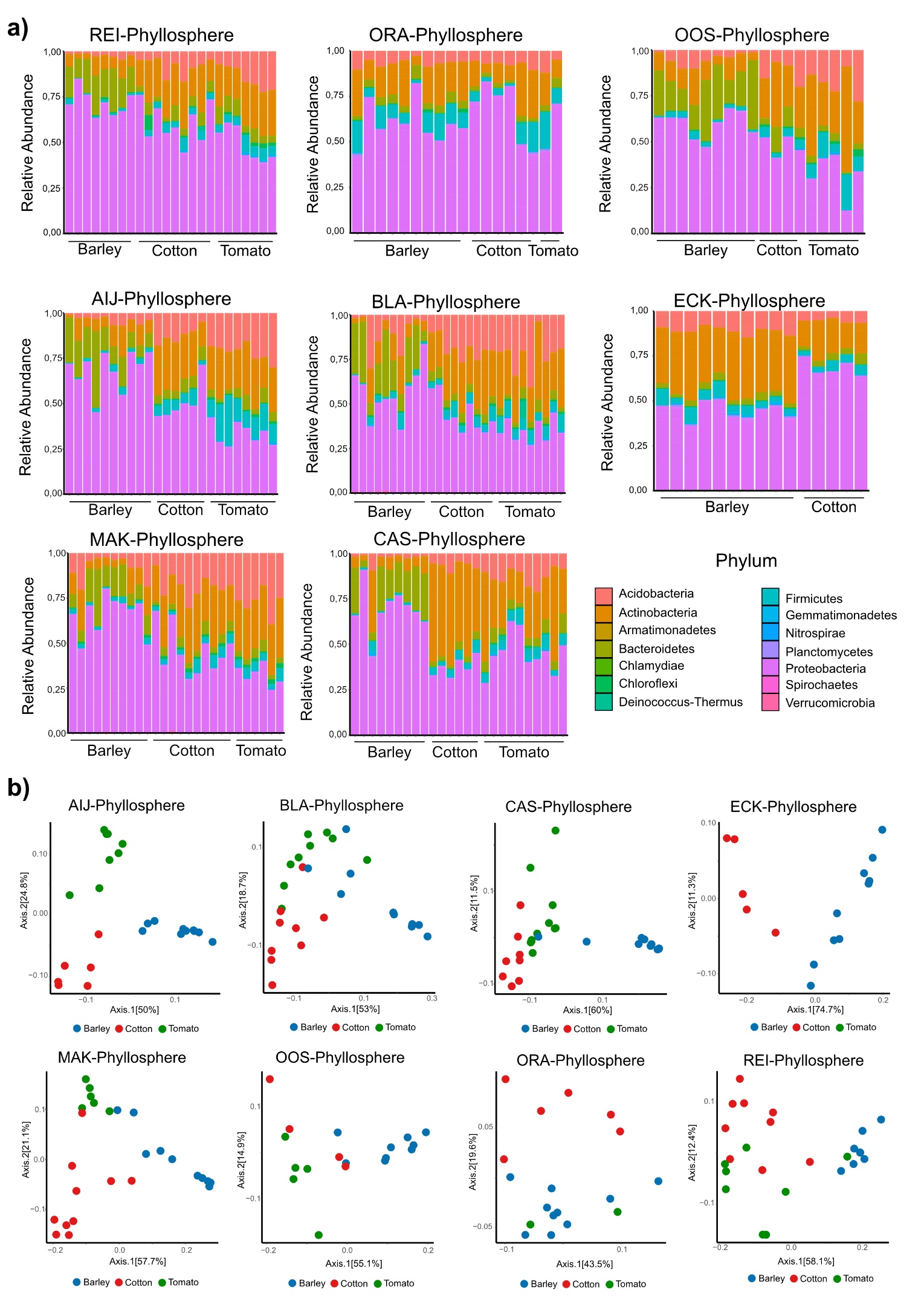

**Supplementary Figure 3. Bacterial phyllosphere microbiota. a)** Relative abundance in percentage of the bacterial phyllosphere microbiota from barley, cotton and tomato plants grown for three weeks on different natural soils. **b)** Principal coordinate analysis (PCoA) based on weighted Unifrac distances of bacterial phyllosphere microbiota from barley, cotton and tomato plants grown on the different natural soils.

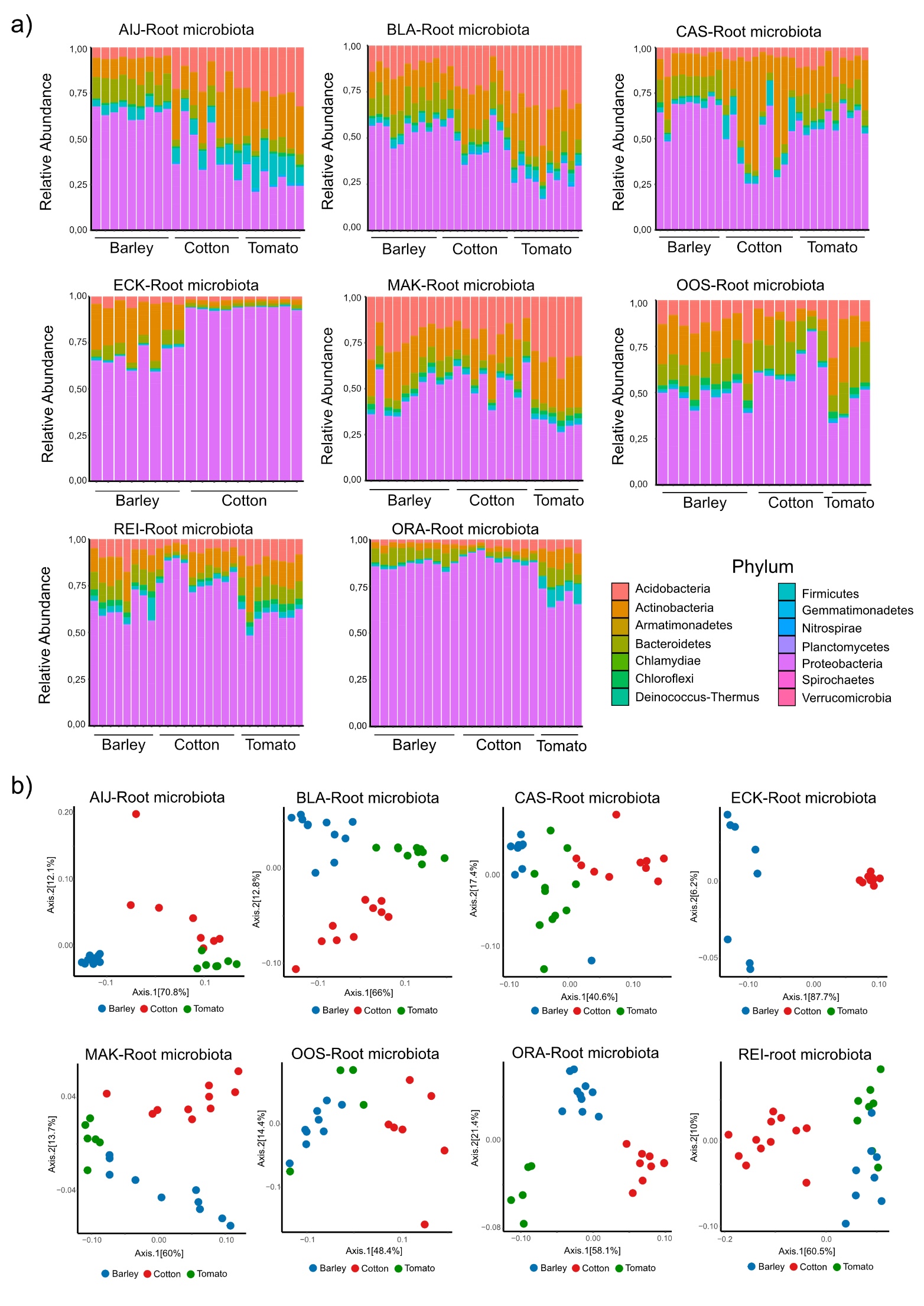

**Supplementary Figure 4. Bacterial root microbiota. a)** Relative abundance in percentage of the bacterial root microbiota of barley, cotton and tomato plants grown for three weeks on different natural soils. **b)** Principal coordinate analysis (PCoA) based on weighted Unifrac distances of the bacterial root microbiota from barley, cotton and tomato plants grown on the different natural soils.

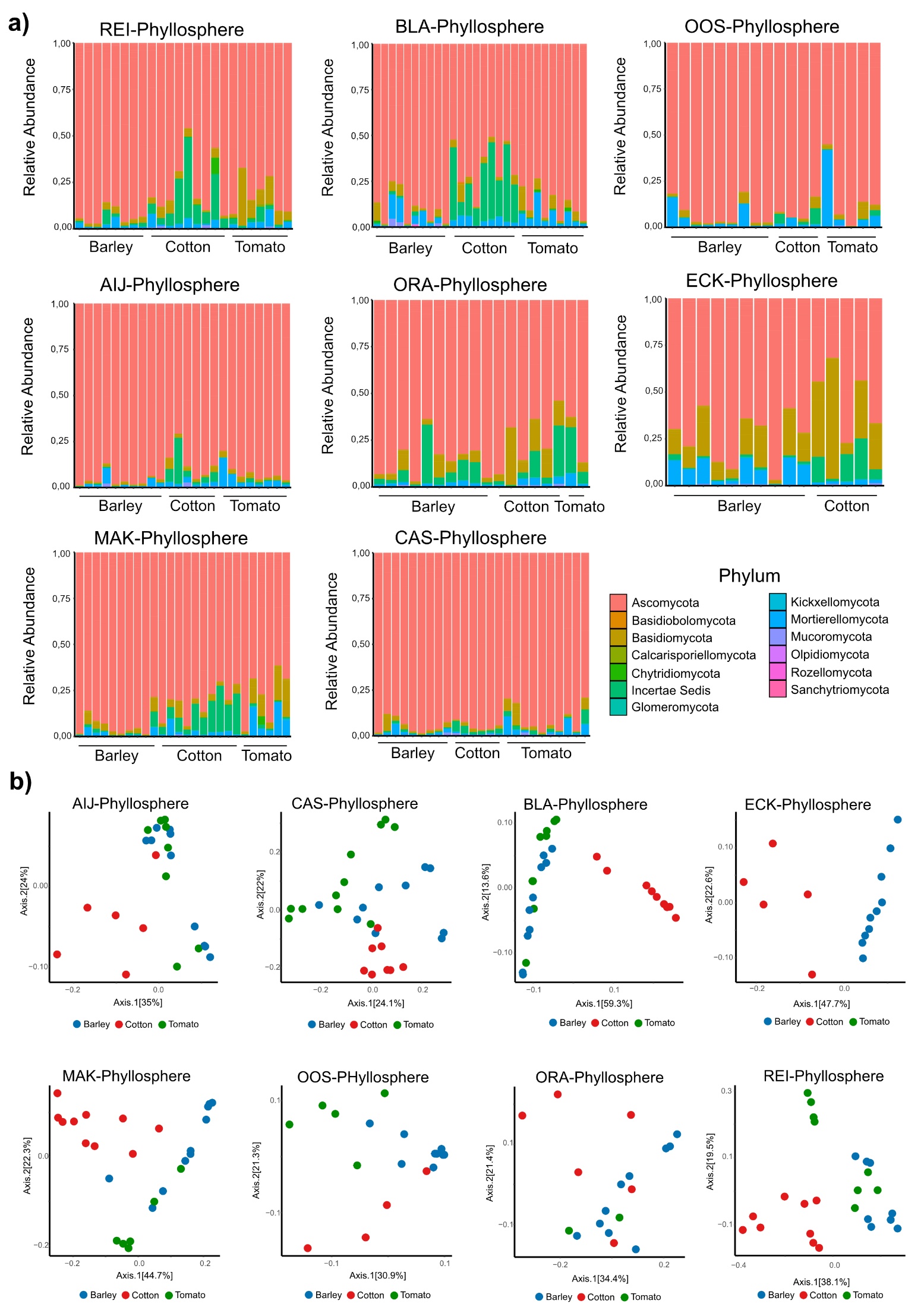

**Supplementary Figure 5. Fungal phyllosphere microbiota. a)** Relative abundance in percentage of the fungal phyllosphere microbiota from barley, cotton and tomato plants grown for three weeks on different natural soils. **b)** Principal coordinate analysis (PCoA) based on weighted Unifrac distances of fungal phyllosphere microbiota from barley, cotton and tomato plants grown on the different natural soils.

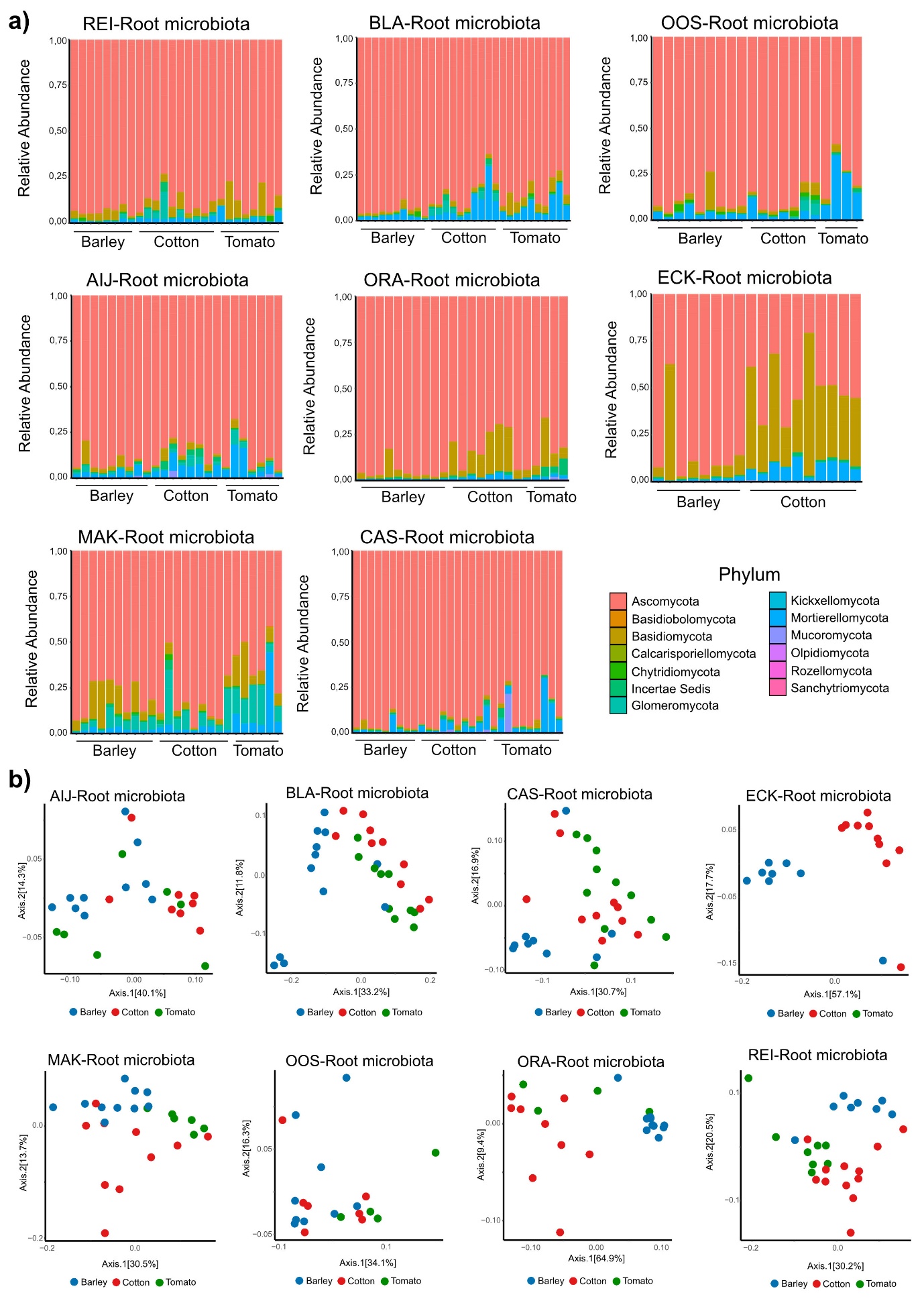

**Supplementary Figure 6. Fungal root microbiota. a)** Relative abundance in percentage of the fungal root microbiota from barley, cotton and tomato plants grown on different natural soils. **b)** Principal coordinate analysis (PCoA) based on weighted Unifrac distances of the fungal root microbiota from barley, cotton and tomato plants grown on the different natural soils.

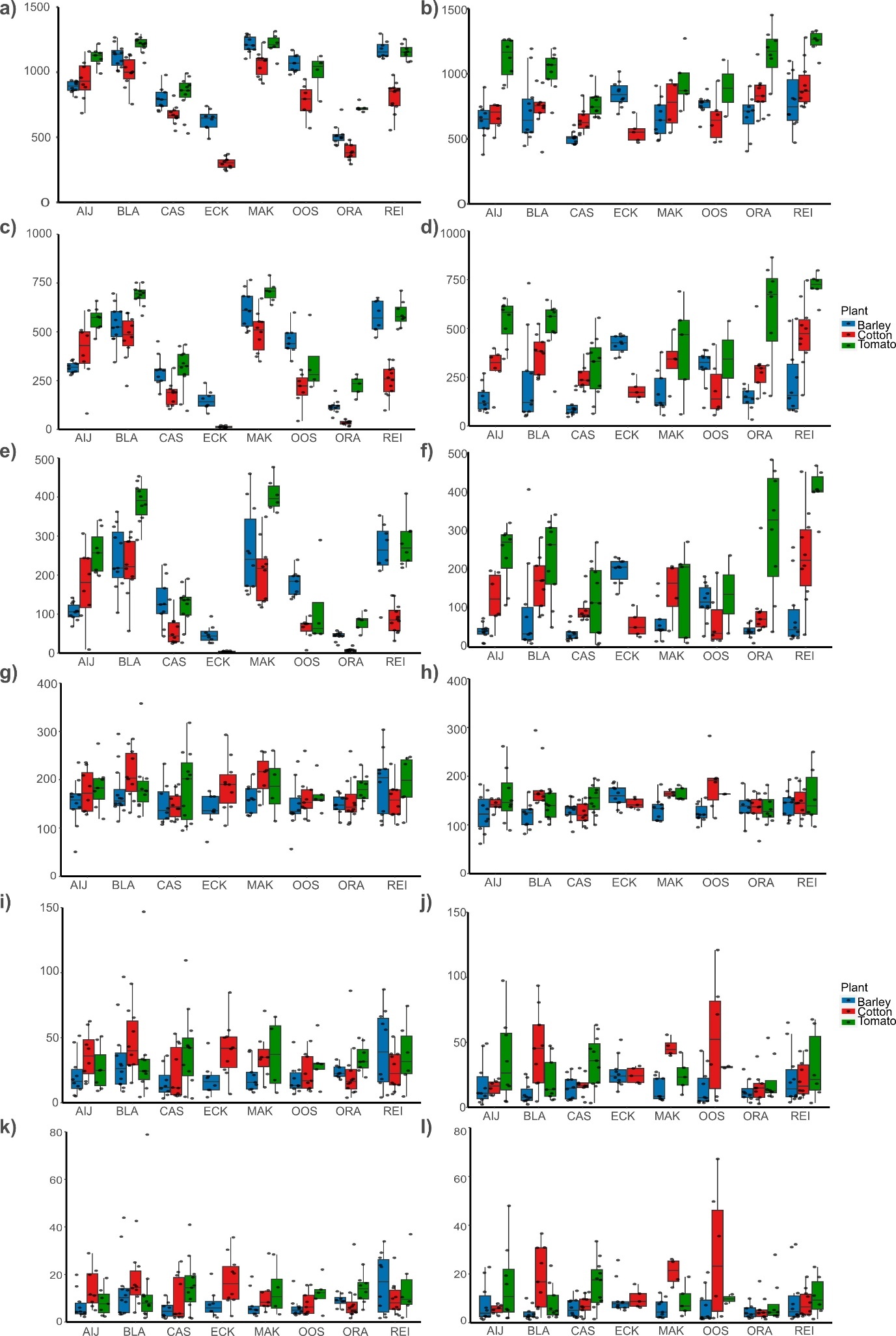

**Supplementary figure 7. α-diversity measurements for bacterial and fungal communities in both the rhizosphere and the phyllosphere of different plant species grown on different natural soils.** a) c) e) correspond to the hill#0, hill#1 and hill#2 of the bacterial rhizosphere microbiota. b), d), f) correspond to the hill#0, hill#1 and hill#2 of the bacterial phyllosphere microbiota. g) i) k) correspond to the hill#0, hill#1 and hill#2 of the fungal rhizosphere microbiota. h), j), l) correspond to the hill#0, hill#1 and hill#2 of the fungal phyllosphere microbiota.

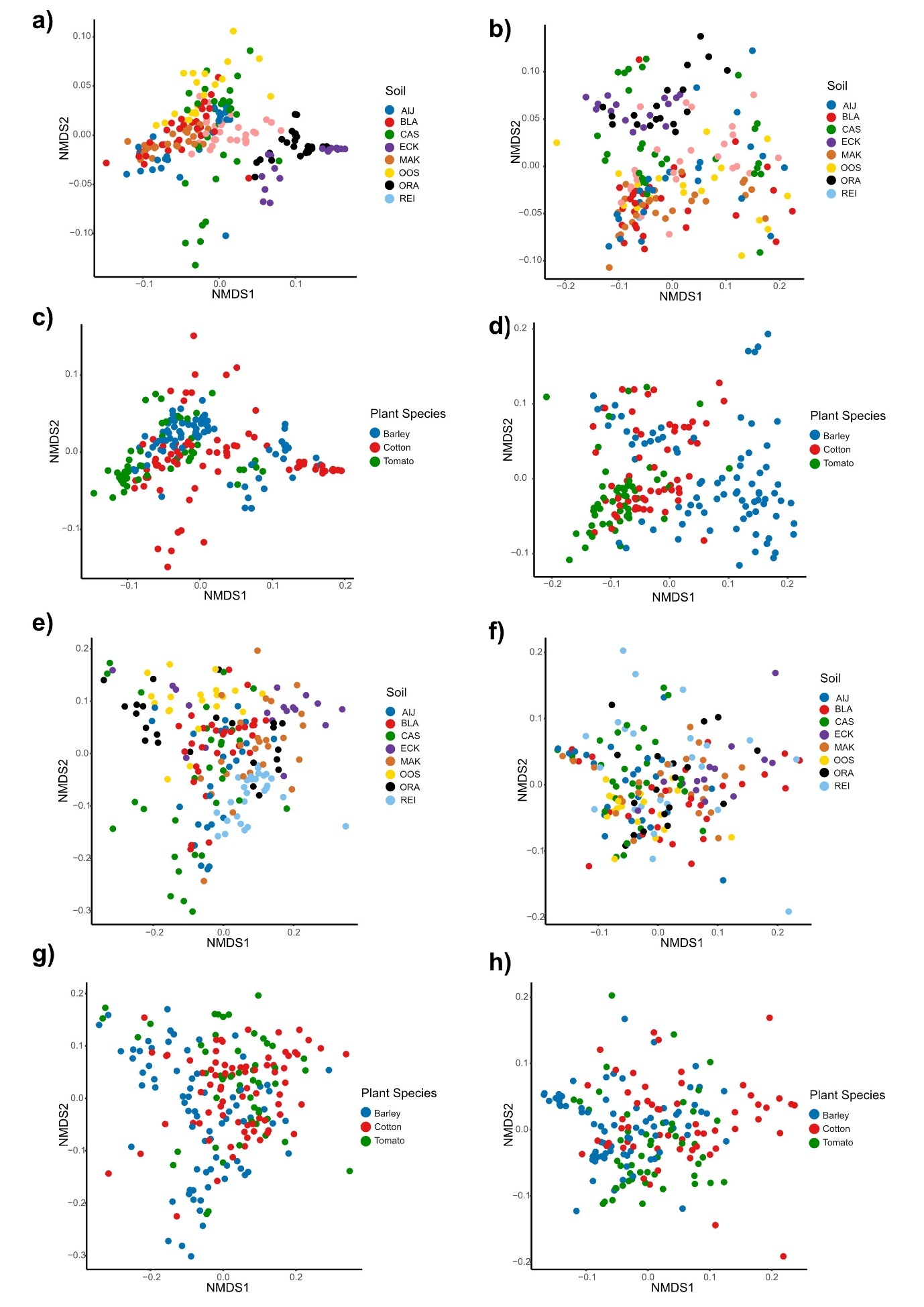

**Supplementary Figure 8. Composition of the root and phyllosphere associated microbiota of barley, cotton and tomato plants grown on the different natural soils. a)** NMDS based on weighted Unifrac distance of root bacteria colored by the type of soil **b)** NMDS based on weighted Unifrac distance of phyllosphere bacteria colored by the type of soil **c)** NMDS based on weighted Unifrac distance of root bacteria colored by plant species **d)** NMDS based on weighted Unifrac distance of phyllosphere bacteria colored by plant species **e)** NMDS based on weighted Unifrac distance of root fungi colored by the type of soil **f)** NMDS based on weighted Unifrac distance of phyllosphere fungi colored by the type of soil **g)** NMDS based on weighted Unifrac distance of root fungi colored by plant species **h)** NMDS based on weighted Unifrac distance of phyllosphere fungi colored by plant species.

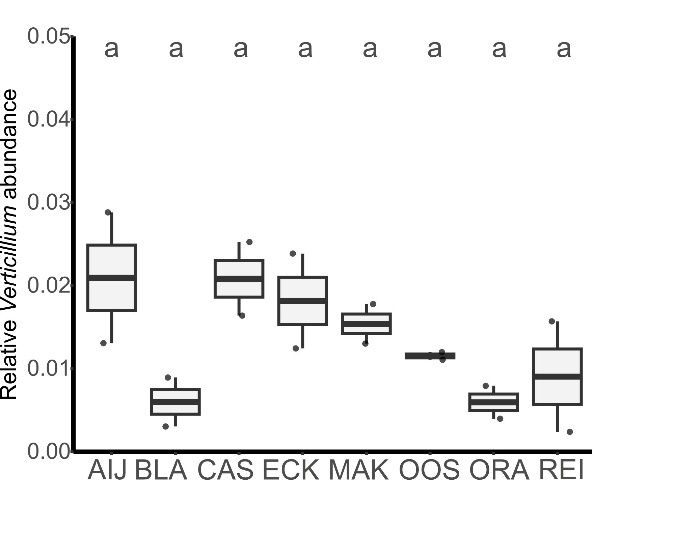

**Supplementary figure 9. There is no significant difference in the relative abundance comparison of *Verticillium* in natural soil.** Different letters represent significant differences (one-way ANOVA and Tukey’s post hoc test; P < 0.05).

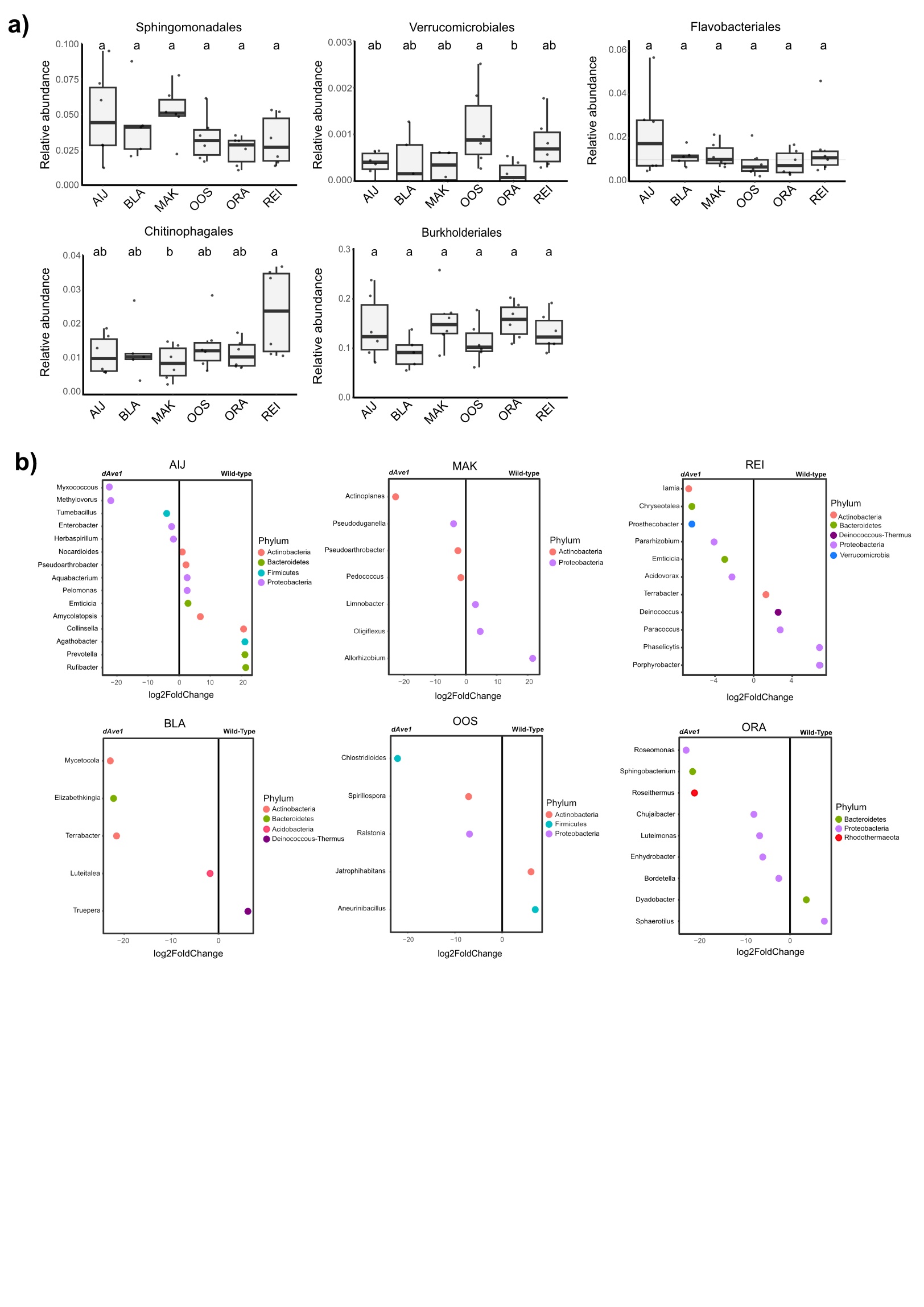

**Supplementary Figure 10. Ave1-mediated microbiota manipulation in the different natural soils. a)** Bacterial families previously reported to be impacted by Ave1 during tomato colonization. Relative abundance of each family in the root microbiota of three-week-old mock-inoculated plants. Different letters indicate statistical differences based on One-Way-Anova (Tukey HSD-Test pval < 0.05). **b)** Differential abundance of bacterial genera between root microbiota of tomato plants inoculated with wild-type *V. dahliae* or an *Ave1* deletion strain (Wald test, adjusted P < 0.05).

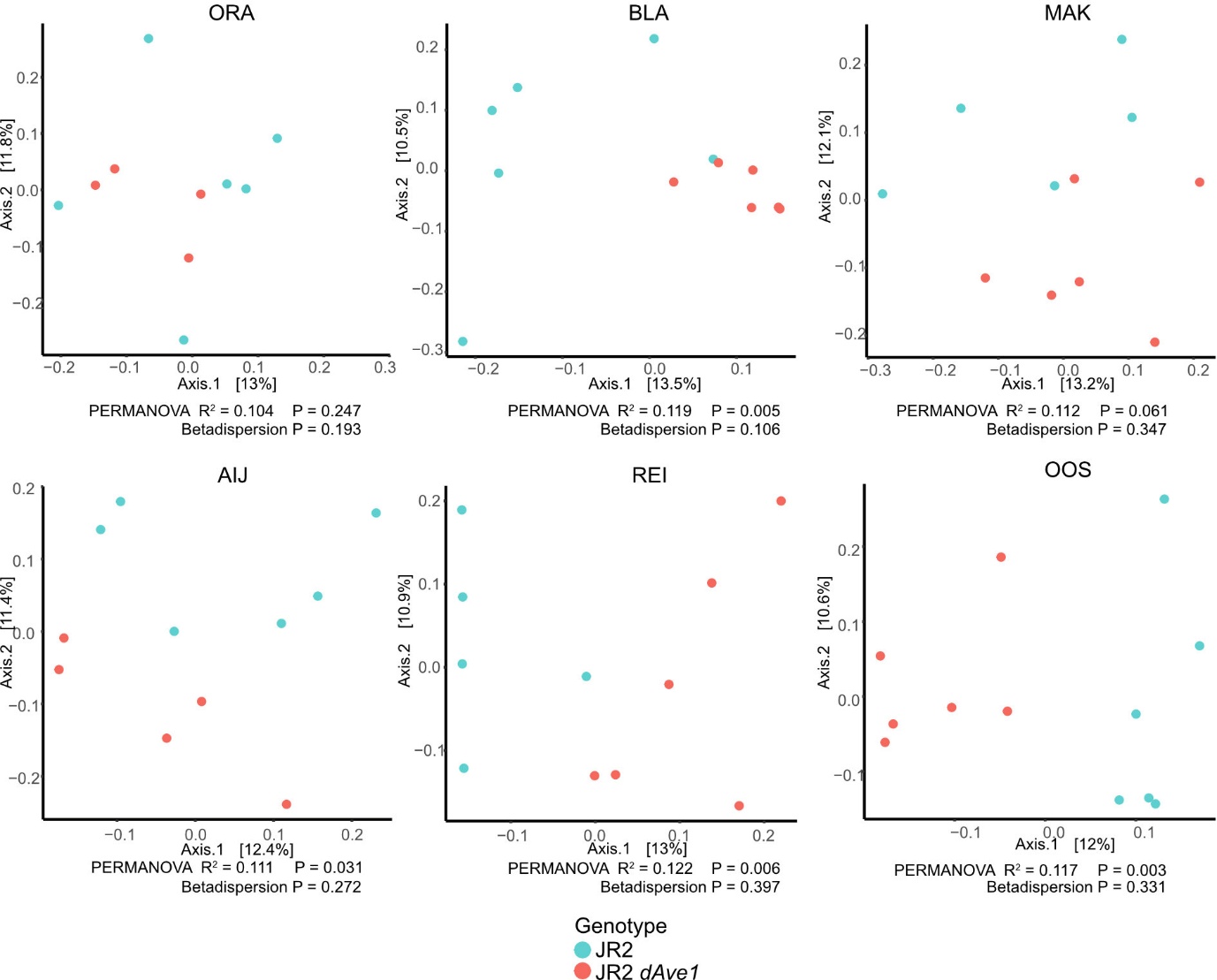

**Supplementary figure 11**. **Microbiota of plants inoculated with wild-type *V. dahliae* or an *Ave1* deletion strain (dAve1) differ significantly.** Principal coordinate analysis (PCoA) based on Unifrac distances of the root microbiota of tomato plants grown on different soils at 14 dpi with wild-type *V. dahliae* (JR2) or an *Ave1* deletion mutant (dAve1). All PERMANOVAs are performed with 9.999 permutations.

**Supplementary Table 1: Locations of the soil collection sites**

| Location | Coordinates | Type of Soil | Abbreviation |
| --- | --- | --- | --- |
| Aijen | 51°34'55.0"N 6°02'27.3"E | River Clay | AIJ |
| De Blauwe Kamer | 51°56'34.4"N 5°37'12.9" | River Clay | BLA |
| Makkum | 53°05'09.8"N 5°26'20.3"E | Sea Clay | MAK |
| Oostvardersplassen | 52°27'50.0"N 5°25'10.8"E | Sea Clay | OOS |
| Eckelrade | 50°47'57.7"N 5°44'42.5"E | Loam | ECK |
| Oranjewoud | 52°57'11.7"N 5°57'45.6"E | Peat | ORA |
| Reijerscamp | 52°00'37.7"N 5°46'25.0"E | Sand | REI |
| De Ginkelse Heide | 52°02'10.7"N 5°43'38.9"E | Sand | GIN |
| Maasduinen | 51°28'34.3"N 6°11'34.9"E | Sand | MAA |
| Cologne | 50°57'27.8"N 6°51'22.4"E | Clay | CAS |
